# Revisiting the *Monascus* genus (Eurotiales, Aspergillaceae): A Multilocus Phylogenetic Approach to Species Delimitation

**DOI:** 10.64898/2026.04.21.719803

**Authors:** Wanping Chen, Shijuan Chen, Lili Jia, Youxiang Zhou, Yanchun Shao, Fusheng Chen

## Abstract

*Monascus* spp. are economically important filamentous fungi that have been utilized in the production of beneficial metabolites such as *Monascus* pigments and monacolin K, as well as in the brewing of some Asian fermented foods. The delimitation of *Monascus* species has traditionally relied on phenotypic traits; however, this morphological classification approach is susceptible to subjective judgments and variations in cultural conditions and also may not necessarily be related to the actual genetic relationship. Consequently, synonymy and misidentification frequently occur in *Monascus* taxonomy, highlighting the urgent need for a convenient and reliable classification system for this genus. In this study, a phylogenetic analysis of 82 representative *Monascus* strains, encompassing all previously recognized species of the genus, was conducted based on the concordance of five gene genealogies (*BenA*, *CaM*, *ITS*, *LSU*, and *RPB2*) to clarify species delimitation and resolve phylogenetic relationships within *Monascus*. The results revealed that the genus *Monascus* is resolved into 11 species, which are clustered into two sections: *Floridani* (including *M. argentinensis*, *M. flavipigmentosus*, *M. floridanus*, *M. lunisporas*, *M. mellicola*, *M. pallens*, and *M. recifensis*) and *Rubri* (including *M. pilosus*, *M. purpureus*, *M. ruber*, and *M. sanguineus*). *M. pilosus* and *M. sanguineus* were reaffirmed as distinct species due to their well-supported and divergent phylogenetic lineages. Additionally, *M. albidulus*, *M*. *anka*, *M. barkeri*, and *M. fumeus* are synonymized with *M. pilosus*, while *M. aurantiacus* and *M. rutilus* are synonyms of *M. purpureus.* Finally, a comprehensive list of accepted *Monascus* species along with their corresponding barcode sequence data is provided.

## Introduction

*Monascus* spp. are filamentous fungi renowned for their rice-fermented product, red mold rice, which has been extensively utilized as fermentation starters, food-grade colorants, and lipid-lowering agents, particularly in East Asia, due to their potent hydrolytic enzymes and beneficial metabolites such as *Monascus* pigments and monacolin K (Chen et al. 2015; Zhu et al. 2019; Feng and Yu 2020; Wu et al. 2021). *Monascus* spp. were first described and nominated by van Tieghem for observing that a cleistothecium was composed of a single spherical ascus in this genus (Mono-ascus) (van Tieghem 1884). Actually, the observation for characterizing *Monascus* is incorrect, however, this genus name has been used ever since. *Monascus* spp can complete their life cycles through asexual or sexual reproduction. The asexual spores (conidia) of *Monascus* spp. are produced either directly from hyphae or laterally on short pedicels either singly or in short chains (Barbosa et al. 2017; Jia et al. 2021). The sexual spores (ascospores) of *Monascus* spp. are formed in a cleistothecium enclosing eight-spored asci. However, when reaching maturity, the ascus wall usually dissolves and ascospores are released into the cleistothecium (Young 1931; Wong and Chien 1986; Patakova 2013). Based on the latest phylogenetic classification, the *Monascus* genus has been placed in the family Aspergillaceae, showing a close relationship with the well-known *Aspergillus* and *Penicillium* (Visagie et al. 2025).

The classification of *Monascus* at species level has traditionally relied on macro- and microscopic features, such as the pigmentation of the cleistothecial walls and conidia, aerial hyphae and growth rates on agar media (Barbosa et al. 2017). Since the introduction of the *Monascus* genus in 1884 (van Tieghem 1884), at least 36 *Monascus* species have been described (**Table 1**), suggesting a large genetic diversity. In the 1930s, Sato isolated numerous *Monascus* strains from fermented products in China, Japan and Korea, classifying them into 13 species (Sato 1936). A systematic review of the genus was conducted by Hawksworth and Pitt (1983). Based on physiological and morphological characteristics, they consolidated a number of *Monascus* species into three: *M. pilosus* (merging *M. pubigerus*, *M. rubropunctatus*, and *M. serorubescens*), *M. ruber* (merging *M. barkeri*, *M. fuliginosus*, *M. heterosporus*, *M. olei*, *M. paxii* and *M. vitreus*), and *M. purpureus* (merging *M. albidus*, *M. albidus* var. *glaber*, *M. anka*, *M. anka* var. *rubellus*, *M. araneosus*, *M. kaoliang*, *M. major*, *M. rubiginosus*, and *M. vini*), while excluding *M. bisporus*, and *M. mucoroides*. In 2004, Li and Guo described four new species *M. albidulus*, *M. rutilus*, *M. fumeus* and *M. aurantiacus*, and provided a retrieval table for 12 *M.* species (including *M. eremophilus*, *M. floridanus*, *M. lunisporas*, *M. pallens*, *M. pilosus*, *M. purpureus*, *M. ruber*, and *M. sanguineus*) by revisiting the genus (Li and Guo 2004). In 2017, Barbosa et al. described three new species *M. flavipigmentosus*, *M. mellicola*, and *M. recifensis* from honey, pollen and the inside of the nest, utilizing a polyphasic approach that combined sequence data, macro- and microscopic characteristics, and extrolites, and transferred the xerophile *M. eremophilus* to *Penicillium eremophilum*.

**Table 1.** List of *Monascus* species described in the literature.

| Species | Remarks | References |
| --- | --- | --- |
| *M. mucoroides* van Tieghem (1884) | The *Monascus* genus was named and two founding species were introduced. Since its discovery, *M. ruber* has consistently been regarded as a distinct species. However, to date, neither the original culture nor other relevant strains of *M. mucoroides* have been isolated or traced. *M. mucoroides* should be excluded as suggested by Hawksworth and Pitt (1983). | (van Tieghem 1884) |
| *M. ruber* van Tieghem (1884) |  |  |
| *M. heterosporus* (Harz) Schröter (1894) | The species was isolated from a raw glycerin solution of 8-10 percent concentration in a candle and soap factory and identified as *Physomyces heterosporus* by Harz (1890), and was later reassigned to *M. heterosporus* by Schröter (1894). In 1983, *M. heterosporus* was regarded as a synonym of *M. ruber* by Hawksworth and Pitt. | (Harz 1890; Schröter 1894; Hawksworth and Pitt 1983) |
| *M. purpureus* Went (1895) | The species was isolated from red mold rice collected in Java. Since then, *M. purpureus* has consistently been regarded as a distinct species. | (Went 1895) |
| *M. barkeri* Dang. (1903) | The species was from the Samsu culture for a traditional rice wine in Southeast Asia and is regarded as a synonym of *M. pilosus* in this study. | (Barker 1903) |
| *M. olei* Piedallu (1910) | The species was found on some oil in cans and on skins from | (Piedallu 1910; |
|  | a tannery in France. In 1983, *M. olei* was regarded as a synonym of *M. ruber* by Hawksworth and Pitt. | Hawksworth and Pitt 1983) |
| *M. paxii* Lingelsh. (1916) | The species was from samples of some *Cluytia* species collected in East Africa. In 1983, *M. paxii* was regarded as a synonym of *M. ruber* by Hawksworth and Pitt. | (Lingelsheim 1916; Hawksworth and Pitt 1983) |
| *M. albidus* (1936) | These 13 species were isolated from fermented products such as red mold rice, wheat leaven, and fermented bean curd in China, Korea, and Japan. However, they were invalidly published and inadequately characterized by modern standards. In 1983, Hawksworth and Pitt reclassified these species into three species *M. pilosus* (merging *M. pubigerus*, *M. rubropunctatus*, and *M. serorubescens*), *M. ruber* (merging *M. fuliginosus*, and *M. vitreus*), and *M. purpureus* (merging *M. albidus*, *M. albidus* var. *glaber*, *M. anka*, *M. anka* var. *rubellus*, *M. araneosus*, *M. major*, and *M. rubiginosus*). *M. pilosus* was regarded as a synonym of *M. ruber* by Barbosa et al. (2017) and is recovered as a distinct species in this study. | (Sato 1936; Hawksworth and Pitt 1983) |
| *M. albidus* var. *glaber* (1936) |  |  |
| *M. anka* Nakaz. & Sato (1936) |  |  |
| *M. anka* var. *rubellus* Sato (1936) |  |  |
| *M. araneosus* Sato (1936) |  |  |
| *M. fuliginosus* Sato (1936) |  |  |
| *M. major* Sato (1936) |  |  |
| *M. pilosus* Sato (1936) |  |  |
| *M. pubigerus* Sato (1936) |  |  |
| *M. rubiginosus* Sato (1936) |  |  |
| *M. rubropunctatus* Sato (1936) |  |  |
| *M. serorubescens* Sato (1936) |  |  |
| *M. vitreus* Sato (1936) |  |  |
| *M. vini* Săvul. & Hulea (1953) | The species was isolated from vino exacidulato in Bucharest, Muntenia, Romania. In 1983, *M. vini* was regarded as a synonym of *M. purpureus* by Hawksworth and Pitt. | (Hawksworth and Pitt 1983) |
| *M. bisporus* (Fraser) Arx (1970) | The species was first described as *Xeromyces bisporus* isolated from liquorice by Fraser in 1954, and later assigned to *M. bisporus* by Arx in 1970. However, it was recovered to *X. bisporus* by Hawksworth and Pitt in 1983. | (Fraser 1954; von Arx 1970; Hawksworth and Pitt 1983) |
| *M. kaoliang* Lin & Iizuka (1975) | The species was isolated from the native starter of kaoliang liquor in Taiwan. In 1983, *M. kaoliang* was regarded as a synonym of *M. purpureus* by Hawksworth and Pitt. | (Lin 1975; Lin and Iizuka 1982; Hawksworth and Pitt 1983) |
| *M. floridanus* Cannon & Barnard (1987) | The species was isolated from roots of *Pinus clausa* in Florida, USA. Since then, *M. floridanus* has consistently been regarded as a distinct species. | (Barnard and Cannon 1987) |
| *M. eremophilus* Hocking & Pitt (1988) | The species was isolated from moldy dried prunes in Sydney, Australia, and was later assigned to *Penicillium eremophilum* by Barbosa et al. (2017). | (Hocking and Pitt 1988; Barbosa et al. 2017) |
| *M. pallens* Cannon, Abdullah & Abbas (1995) | They were isolated from surface sediments of the Shatt-al-Arab river and its creeks near Basrah City, Iraq. Since then, *M. pallens* has consistently been regarded as a distinct species. *M. sanguineus* was regarded as a synonym of *M. purpureus* by Barbosa et al. (2017), and is recovered as a distinct species in this study. | (Cannon et al. 1995; Barbosa et al. 2017) |
| *M. sanguineus* Cannon, Abdullah & Abbas (1995) |  |  |
| *M. lunisporas* Udagawa & Baba (1998) | The species was isolated from mouldy feeds in Tokyo, Japan. Since then, *M. lunisporas* has consistently been regarded as a distinct species. | (Udagawa and Baba 1998) |
| *M. argentinensis* Stchigel & Guarro (2004) | The species was isolated from soil samples collected around Tafí del Valle, Tucumán province, Argentina. | (Stchigel et al. 2004) |
| *M. albidulus* Li & Guo (2004) | The species was erected to replace the invalid *Monascus albidus* K. Sato (1936) and is regarded as a synonym of *M. pilosus* in this study. | (Li 1982; Li and Guo 2004) |
| *M. aurantiacus* Li ex Li & Guo (2004) | The species was isolated from a Daqu of Wangjiang brewery, Anhui, China in 1982, and is regarded as a synonym of *M. purpureus* in this study. |  |
| *M. fumeus* Li & Guo (2004) | The species was erected to replace the invalid *M. fuliginosus* Sato (1936) and is regarded as a synonym of *M. pilosus* in this study. |  |
| *M. rutilus* Li & Guo (2004) | The species was erected to replace the invalid *M. anka* Sato (1936) and is regarded as a synonym of *M. purpureus* in this study. |  |
| *M. flavipigmentosus* Barbosa, Souza-Motta, Oliveira & Houbraken (2017) | They were isolated from honey, pollen and inside nests of *Melipona scutellaris* bees in Recife, Pernambuco, Brazil, and have been regarded as distinct species to date. | (Barbosa et al. 2017) |
| *M. mellicola* Barbosa, Souza-Motta, Oliveira & Houbraken (2017) |  |  |
| *M. recifensis* Barbosa, Souza-Motta, Oliveira & Houbraken (2017) |  |  |
| *M. purpureus* var. *aurantiacus* He & Liu (2018) | *Monascus aurantiacus* was regarded as a variant of *M. purpureus* by He et al. (2018) and is regarded as a synonym of *M. purpureus* in this study. | (He et al. 2018) |
| *M. purpureus* var. *rutilus* He & Liu (2018) | *Monascus rutilus* was regarded as a variant of *M. purpureus* by He et al. (2018) and is regarded as a synonym of *M. purpureus* in this study. |  |
| *M. ruber* var. *albidulus* He & Liu (2018) | *Monascus albidulus* was regarded as a variant of *M. ruber* by He et al. (2018) and is regarded as a synonym of *M. pilosus* in this study. |  |
| *M. ruber* x *sanguineus* He & Liu (2018) | *M. barkeri* was regarded as a nothospecies of *M. ruber* and *M. sanguineus* by He et al. (2018) and is regarded as a synonym of *M. pilosus* in this study |  |

Meanwhile, various genetic markers and information have been used to delineate the boundaries of *Monascus* species, which are unaffected by the variations in cultural conditions on morphological and physiological traits (Shao et al. 2014). The random amplified polymorphic DNA (RAPD) marker was initially employed to investigate the genetic relationship among 25 *Monascus* isolates, revealing four distinct *Monascus* genetic lineages (Lakrod et al. 2000). Similar efforts were also made using genetic markers such as the D1/D2 regions of the large subunit (*LSU*) rRNA genes, the internal transcribed spacer (*ITS*), partial β-tubulin (*BenA*) genes, 18S rDNA, inter-simple sequence repeats, and sequence-related amplification polymorphism (Park and Jong 2003; Park et al. 2004; Suharna et al. 2005; Shinzato et al. 2009; Shao et al. 2011; Lv et al. 2012; Wei et al. 2012; Li et al. 2017; Yang et al. 2017; Li et al. 2021; Zhou et al. 2023). In 2017, by integrating morphological characteristics, *ITS*, *LSU*, *BenA*, calmodulin (*CaM*) and RNA polymerase II second largest subunit sequences (*RPB2*) along with extrolite data, Barbosa et al. proposed that 1) the genus *Monascus* is divided into two new sections, *Floridani* and *Rubri*; 2) *M. eremophilus* is reclassified as *Penicillium eremophilum*; 3) *M. pilosus* and *M. sanguineus* are synonyms of *M. ruber* and *M. purpureus*, respectively (Barbosa et al. 2017). In 2018, based on the phylogenetic analysis of *ITS*, *LSU* and *pksKS* sequences alongside morphological analysis, He et al. proposed that 1) *M. aurantiacus* is a variant of *M. purpureus*; 2) *M. rutilus* is a variant of *M. purpureus*; 3) *M. albidulus* is a variant of *M. ruber*; 4) *M. barkeri* is a nothospecies of *M. ruber* and *M. sanguineus*; 5) *M. fumeus* is a synonym of *M. ruber*; 6) *M. sanguineus* is not a synonym of *M. purpureus* (He et al. 2018). In 2020, the phylogenetic analysis of various genetic markers among *M.* strains suggested that *M. sanguineus* may be a natural nothospecies, with its haplotype A closely related to *M. ruber*, while haplotype B may derive from an unknown *Monascus* species (He et al. 2020). In 2022, based on the phylogenetic analysis of *BenA* genes and genome-wide features such as repertoires of secondary metabolite biosynthesis gene clusters, Liu et al. proposed that 1) the genus *Monascus* can be genetically delimited into three sections; 2) *M. sanguineus* should be treated as a separate species from *M. purpureus* (Liu et al. 2022). In 2023, the phylogenetic analysis of 15 *Monascus* strains at the genome level revealed a high homology between *M. pilosus* and *M. ruber*, along with their distant relationship with *M. purpureus*, classifying them into two distinctly evolutionary clades: the *M. purpureus* clade and the *M. pilosus*-*M. ruber* clade (Zhang et al. 2023).

Establishing convenient and reliable classification systems are of great importance for both *Monascus* research and application. In this study, the phylogenetic relationships of 82 representative *Monascus* strains, encompassing all currently recognized species within the genus, were elucidated by analyzing the sequences of the genetic markers *BenA*, *CaM*, *ITS*, *LSU* and *RPB2*. Based on these findings, an updated list of accepted *Monascus* species in the genus together with their corresponding barcode sequence data is provided herein.

## Materials and methods

### *Monascus* reference strains

A total of 82 *Monascus* reference strains from various habitats, regions or countries, encompassing all 17 previously recognized species in the genus, were utilized in this study and their information is provided in **Table 2**. They were mostly sourced from the CABI culture collection (formerly International Mycological Institute, IMI) in the UK, the Korean Collection for Type Cultures (KCTC) and the Korean Agricultural Culture Collection (KACC) in South Korea, the CBS-KNAW culture collection in the Netherlands, Osaka University (OUT) in Japan, the American Type Culture Collection (ATCC) and the ARS Culture Collection (NRRL) in USA, the Micoteca URM of Federal University of Pernambuco in Brazil, the China Center of Industrial Culture Collection (CICC), the Chinese Academy of Sciences (AS), and the China General Microbiological Culture Collection Center (CGMCC).

**Table 2.**
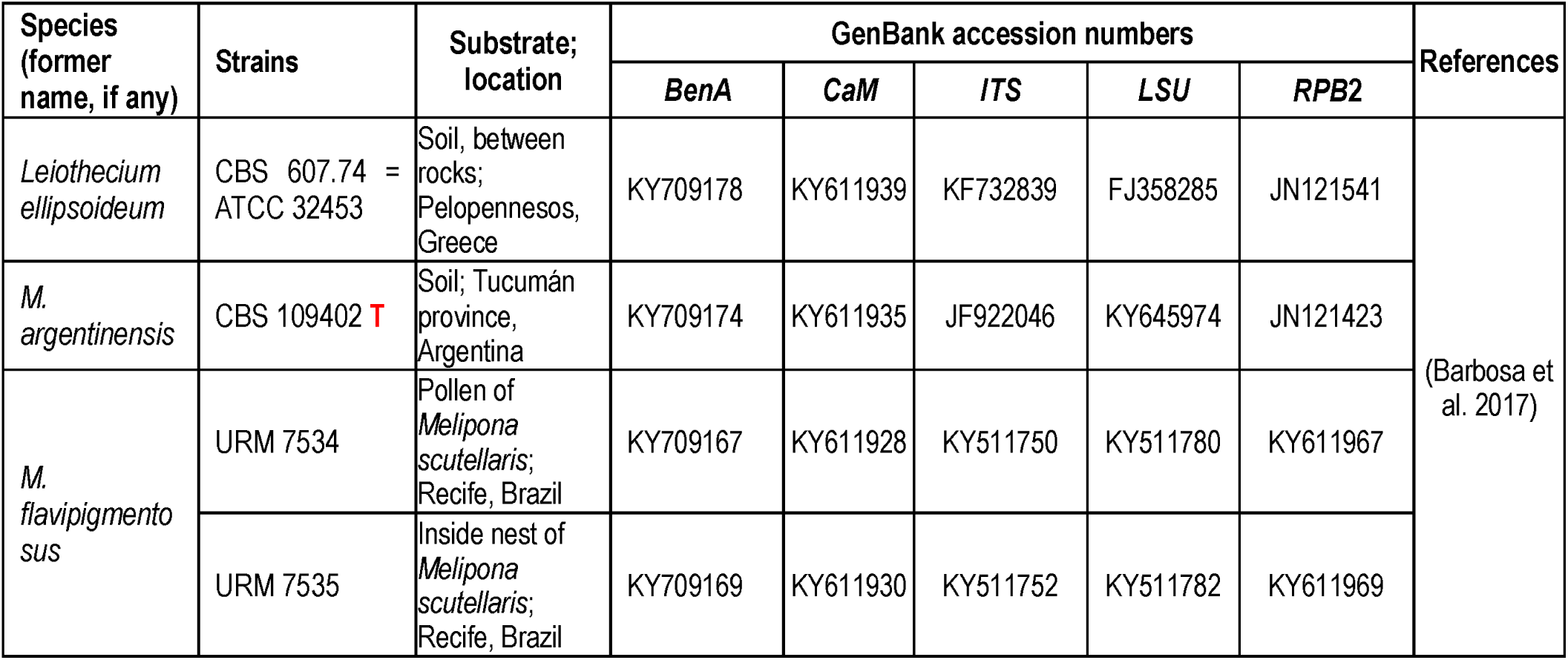
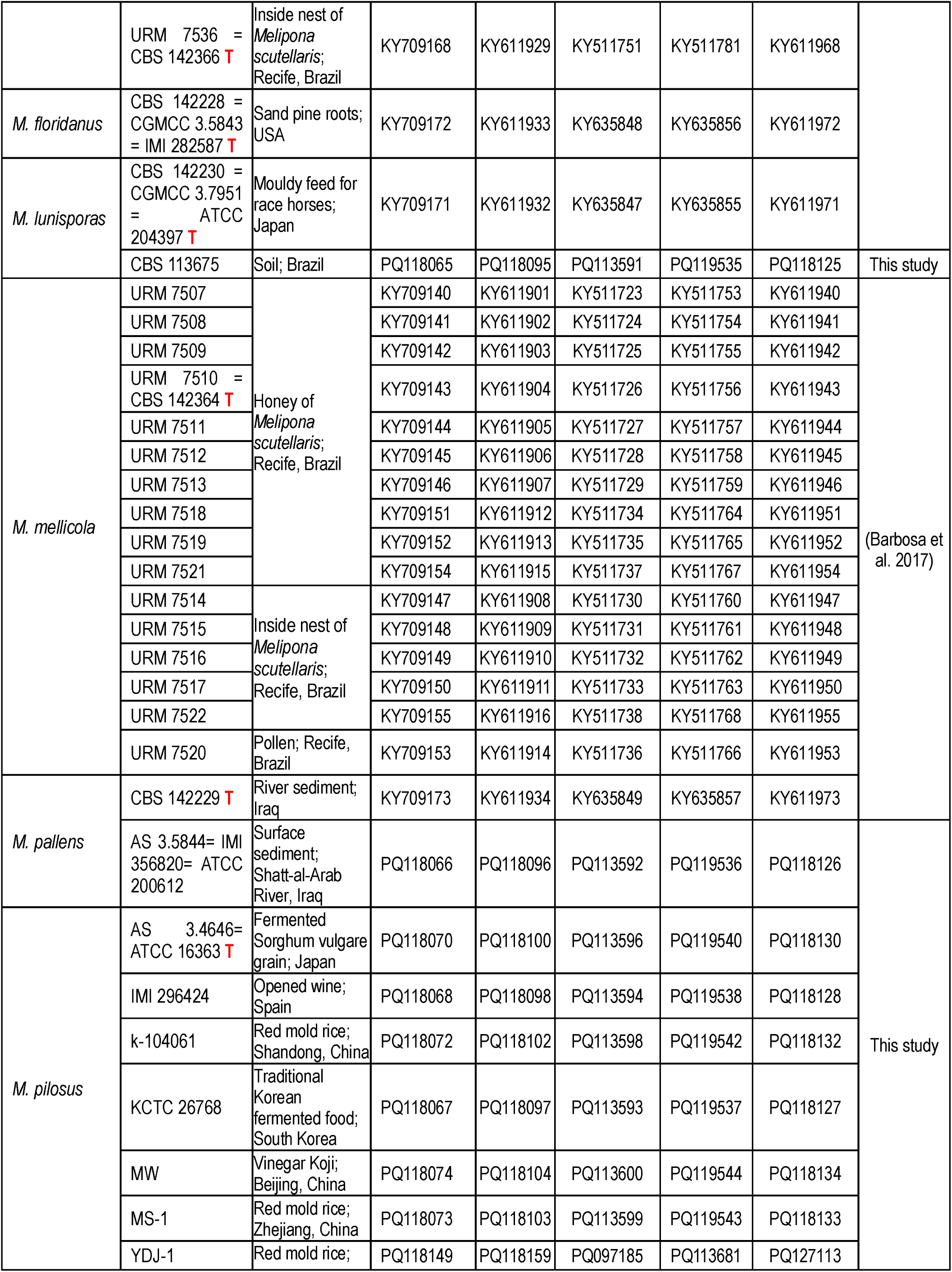
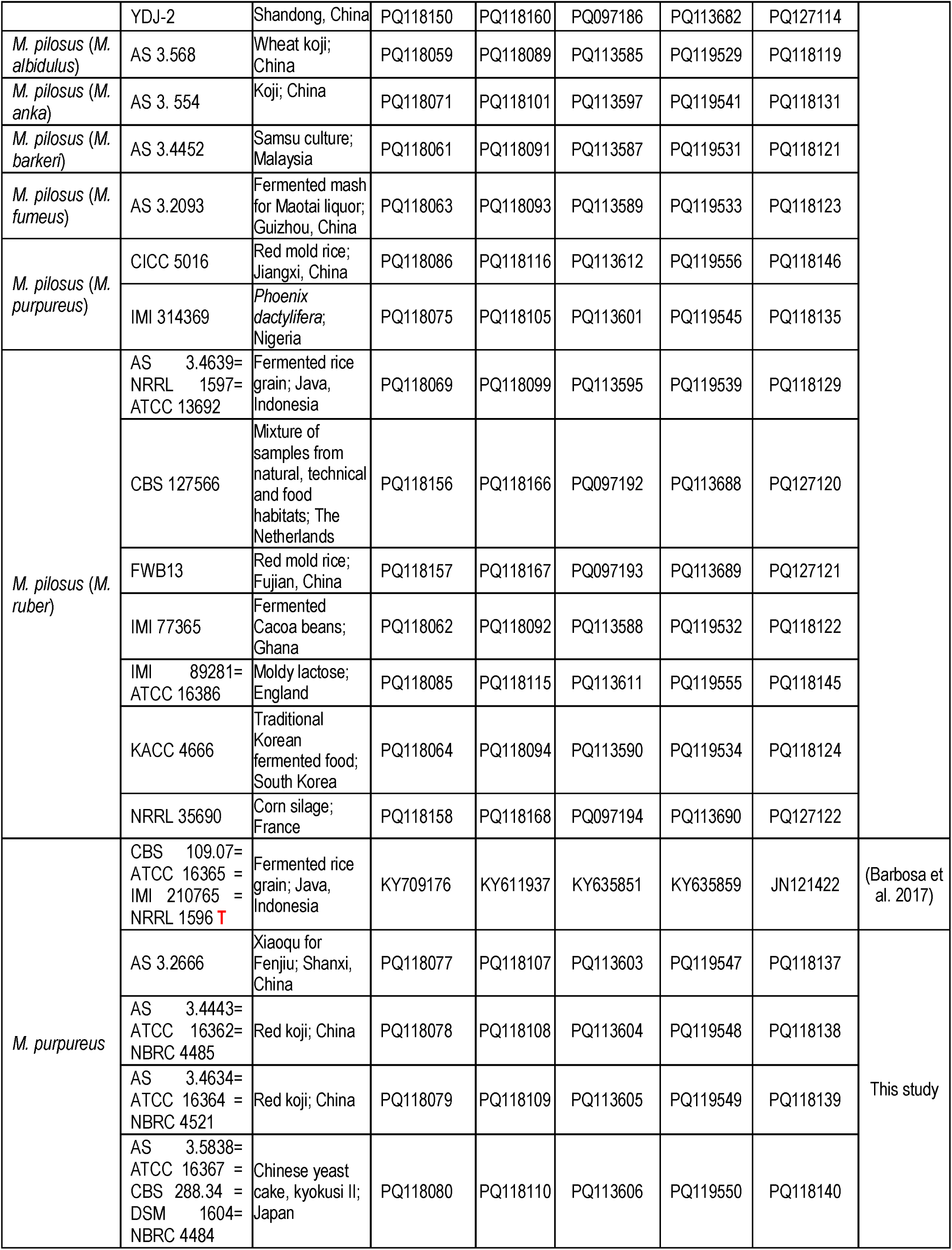
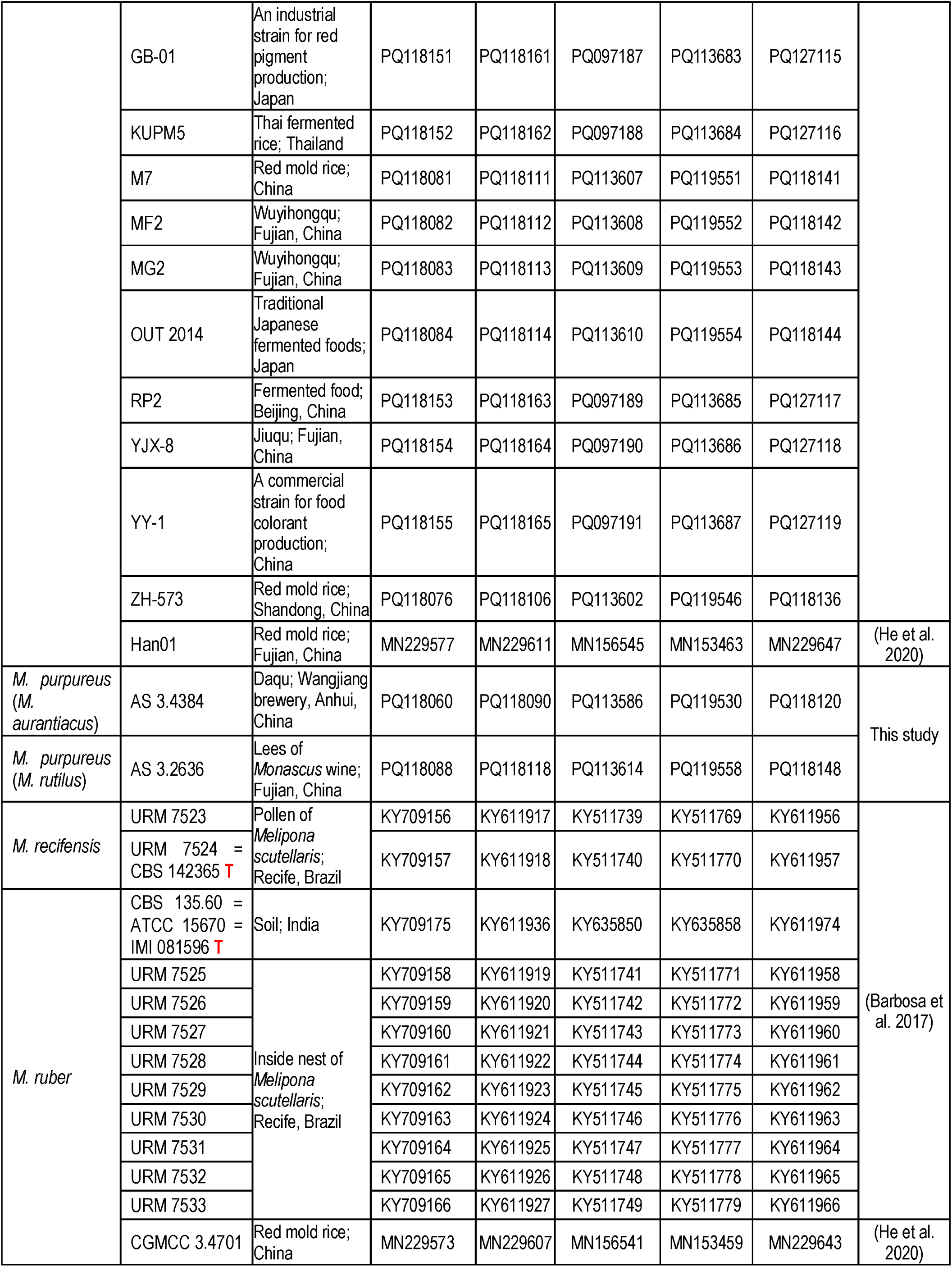
*Monascus* strains and their sequences used in this study.

### DNA extraction, PCR and sequencing

The sequence information of the *ITS*, *LSU*, *BenA*, *CaM* and *RPB2* genes was extracted from the studied *Monascus* strains. Briefly, the genomic DNA of the *Monascus* strains was extracted using cetyltrimethylammonium bromide buffer, following the previous description (Shao et al. 2009) for the amplification of *ITS*, *LSU*, *BenA*, *CaM* and *RPB2* gene sequences by PCR. The primers used for PCR are listed in **Table 3**. The size of the PCR products was verified by agarose gel electrophoresis, and the gene information was obtained through sequencing. Newly generated sequences were deposited in the GenBank.

**Table 3.**
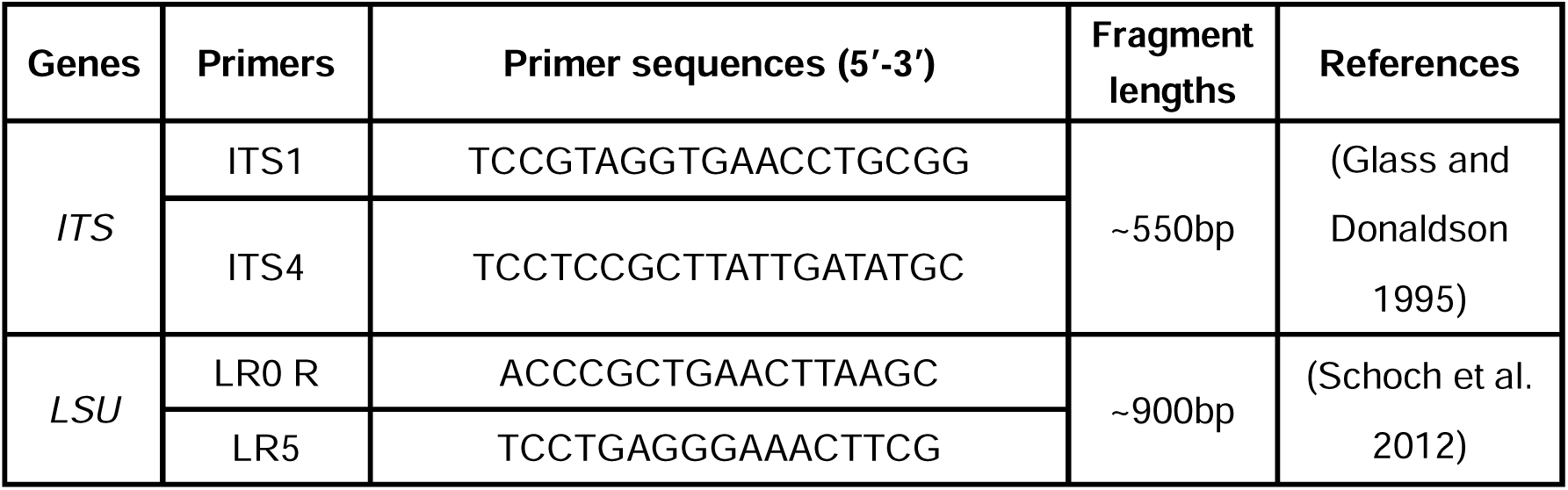
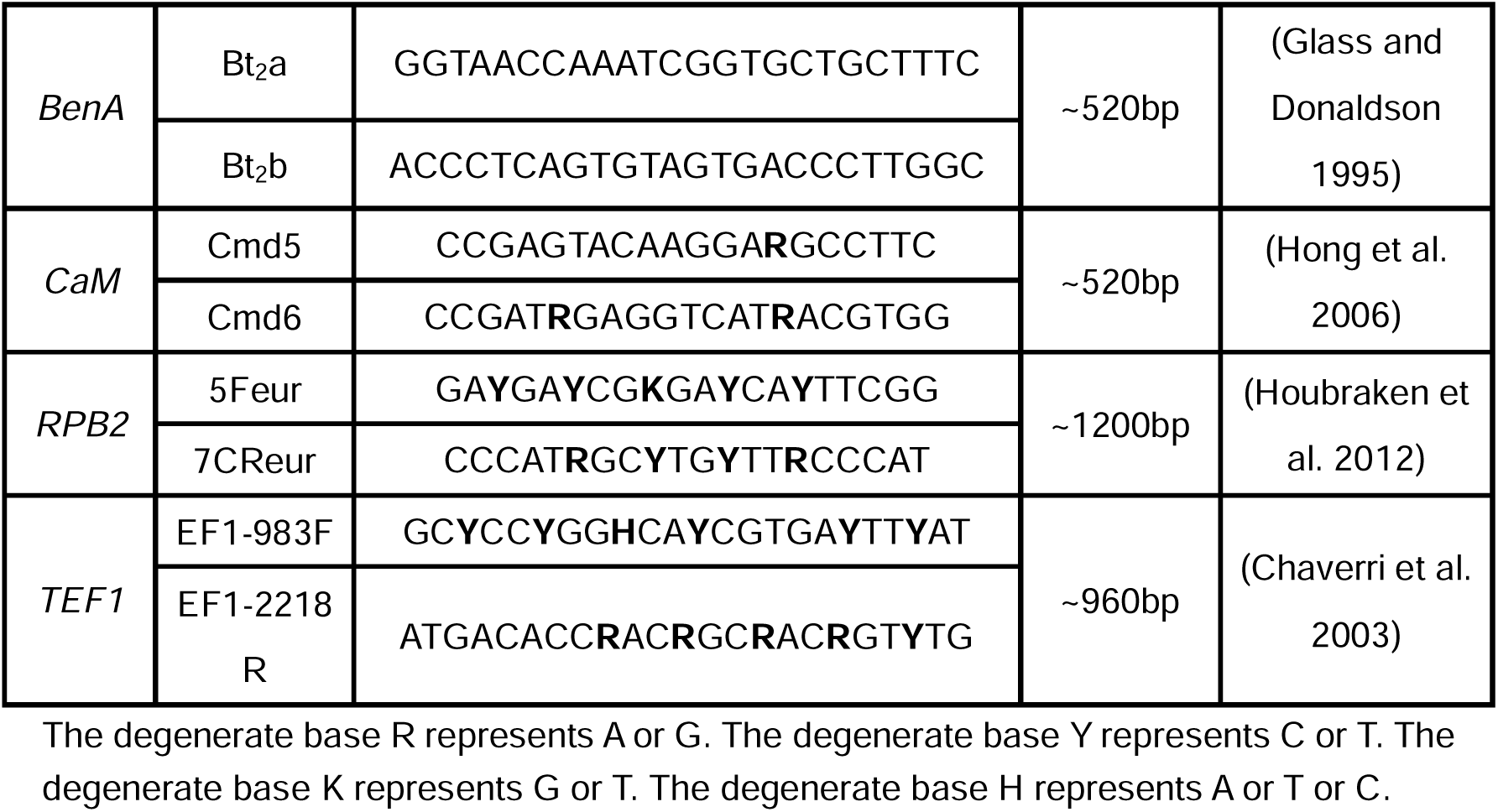
Primers for gene amplification and fragment length.

### Phylogenetic analyses

The DNA sequences of *BenA*, *CaM*, *ITS*, *LSU* and *RPB2* from representative *Monascus* strains were utilized for phylogenetic analyses, and their information is listed in **Table 2**. As previously reported, the corresponding gene regions of *Leiothecium ellipsoideum* strain CBS 607.74 (Aspergillaceae, Eurotiales) were used as out-groups (Barbosa et al. 2017). The sequences were aligned using MAFFT v7.505 (Katoh and Standley 2013). Individual alignments were concatenated by using PhyloSuite v1.2.3 (Zhang et al. 2020). The ModelFinder v2.2.0 (Kalyaanamoorthy et al. 2017) on IQ-TREE software v1.6.12 (Nguyen et al. 2015) was employed to find the best-fitting model for each alignment according to the Bayesian Information Criterion. Phylogenetic trees were constructed using the maximum likelihood methods implemented in IQ-TREE software (Minh et al. 2020b). Branch support values were measured using ultrafast bootstraps from 1,000 replicates, SH-like approximate likelihood ratio tests from 1,000 replicates, and approximate Bayes tests (Hoang et al. 2018; Minh et al. 2020). Trees were visualized and edited in iTOL v6 (Letunic and Bork 2024).

## Results

### Molecular typing of *Monascus* strains

82 representative *Monascus* strains covering the formerly accepted 17 *Monascus* species were collected, and their phylogenetic relationships were investigated using individual or concatenated five gene data set (*BenA*, *CaM*, *ITS*, *LSU* and *RPB2*) (**Table 2**).

The single gene phylogenetic trees for the *BenA*, *CaM*, *ITS*, *LSU* and *RPB2* gene regions of 82 *Monascus* strains are illustrated in **Fig. 1**. The aligned data sets of *BenA*, *CaM*, *ITS*, *LSU* and *RPB2* are 484 bp, 524 bp, 641 bp, 883 bp, and 1029 bp, respectively, including alignment gaps. The most optimal nucleotide substitution models for *BenA*, *CaM*, *ITS*, *LSU* and *RPB2* are HKY+F+G4, K2P+G4, TN+F+G4, TPM2u+F+I, and K2P+G4, respectively, and the trees were constructed under their best optimal models. Overall, a similar topology in section *Floridani* was observed across the five single gene phylogenetic trees. However, the single gene phylogenetic trees lacked sufficient resolution to differentiate *Monascus* species in section *Rubri*. The species *M. pilosus*, *M. ruber* and *M. sanguineus* were closely related or mixed in the five single trees, particularly in the *ITS* and *LSU* phylograms.

**Figure 1.**
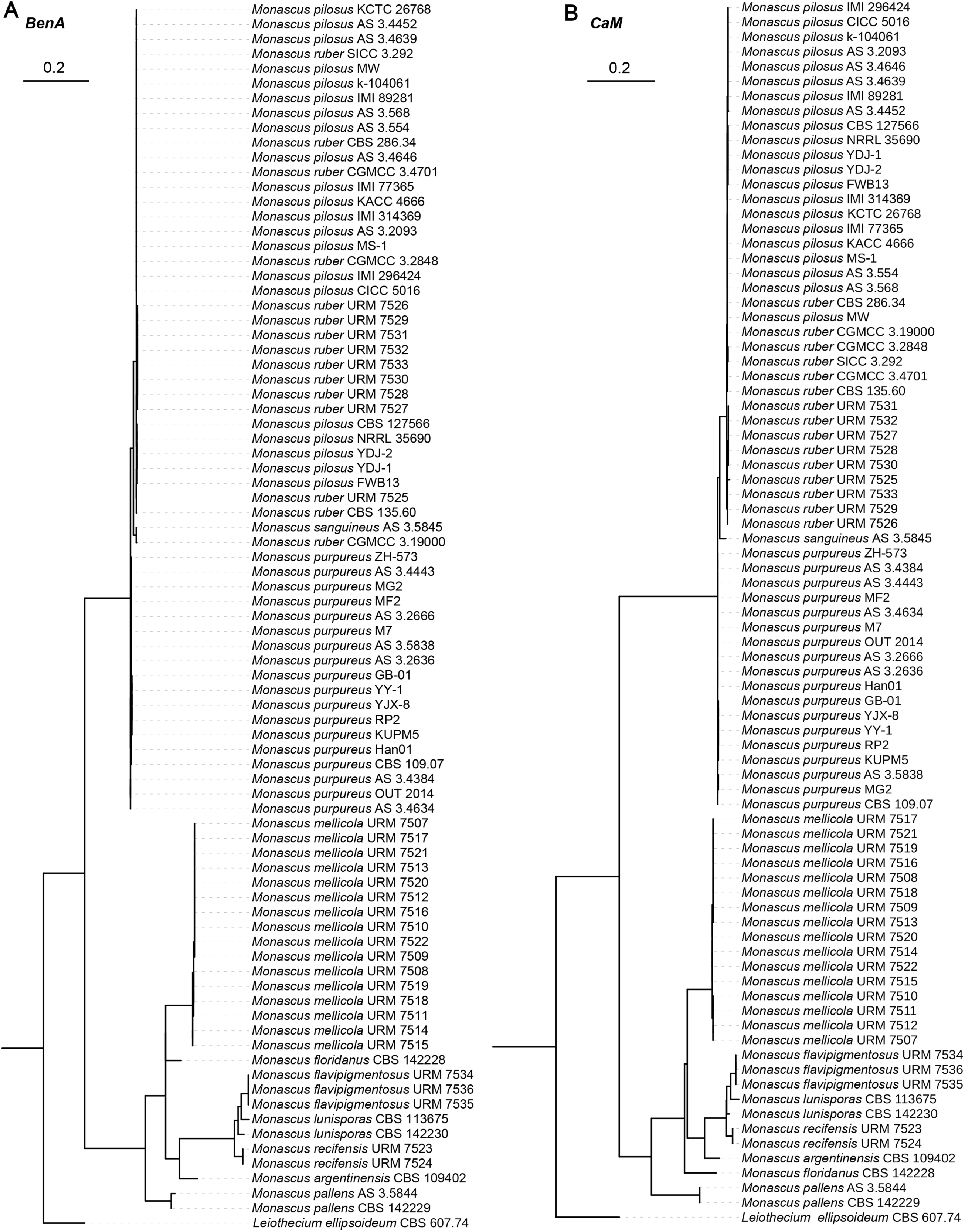
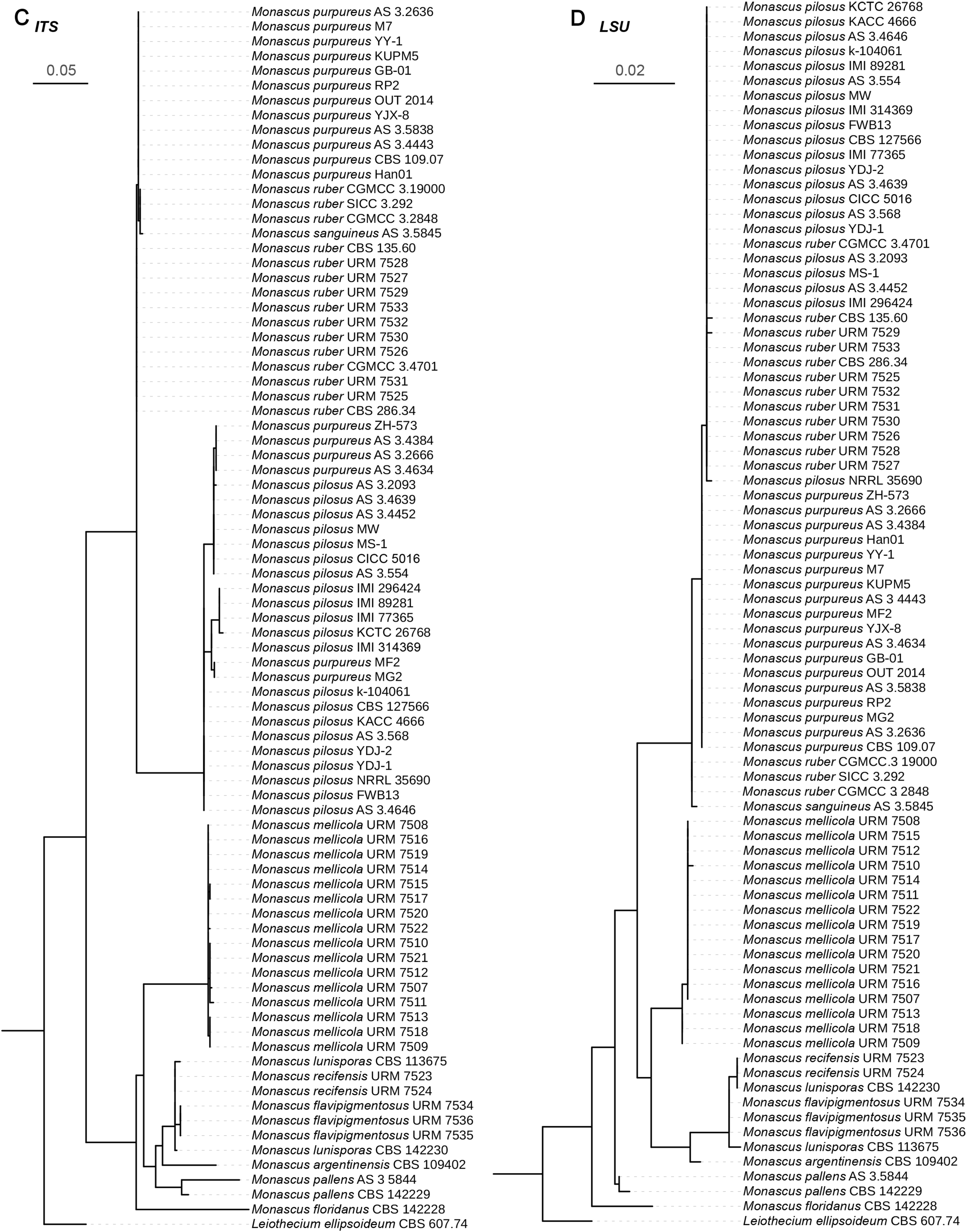
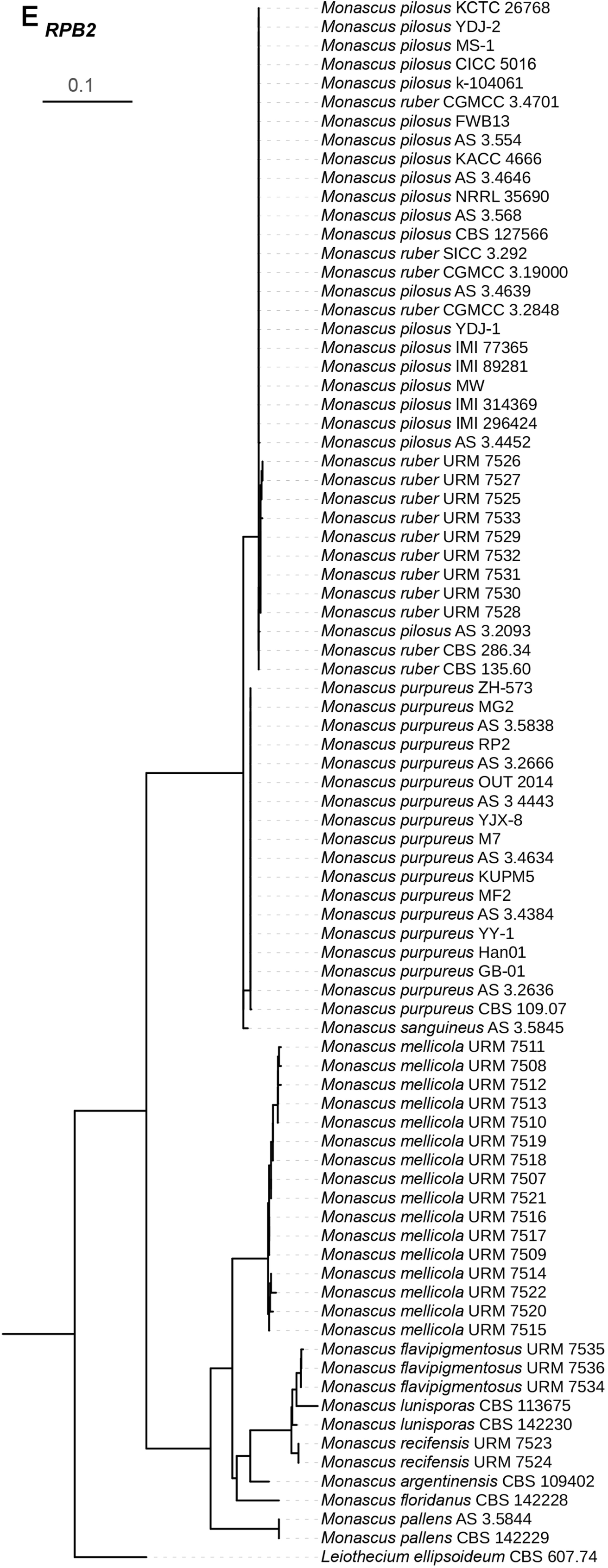
Single gene phylogenetic trees of the *BenA*, *CaM*, *ITS*, *LSU*, and *RPB2* gene regions of *Monascus* strains. Panels A-E correspond to the trees of *BenA*, *CaM*, *ITS*, *LSU* and *RPB2*, respectively.

Therefore, the five gene datasets were concatenated to approach a higher resolution tree for species recognition (**Fig. 2**). The tree was deposited in the TreeBASE under accession number 31619. As previously reported (Barbosa et al. 2017), the analysis revealed two well-supported sections, *Floridani* and *Rubri*, in the *Monascus* genus. In section *Floridani*, there are seven well-supported and distinct species (*M. argentinensis*, *M. flavipigmentosus*, *M. floridanus*, *M. lunisporas*, *M. mellicola*, *M. pallens*, and *M. recifensis*). These seven *Monascus* species were all isolated from natural habitats, rather than from traditional fermented products. Except for *M. flavipigmentosus*, *M. lunisporas*, and *M. recifensis*, which are resolved as close relatives in a distinct, well-supported clade, the other four species exhibit clear genetic differences.

**Figure 2.**
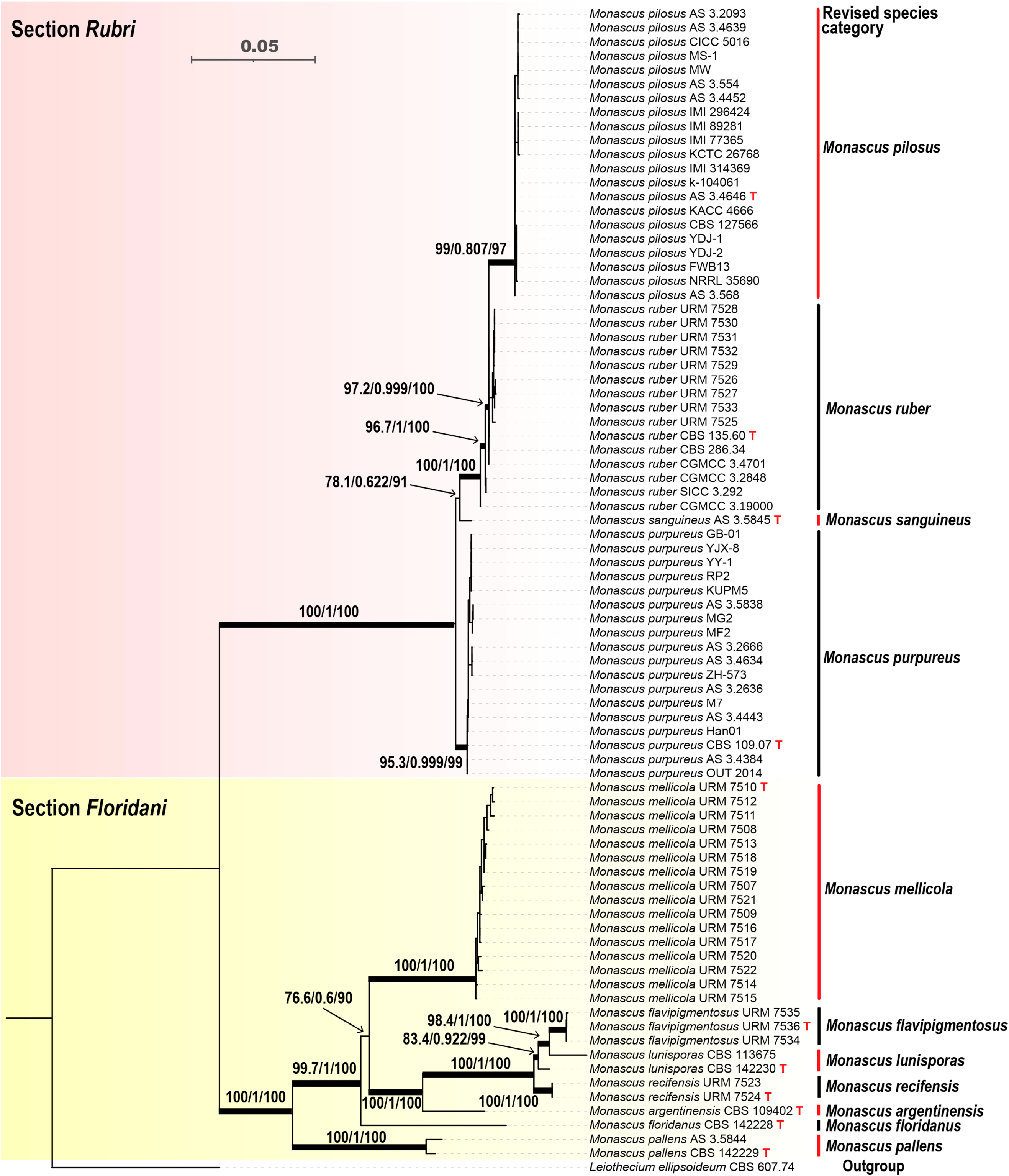
Concatenated phylogeny of the CaM, BenA, LSU, ITS and RPB2 gene regions illustrating the relationship within *Monascus*. Numbers on the branches indicate SH-aLRT support (%) / aBayes support / ultrafast bootstrap support (%) and those branches with values of >95 % bootstrap are thickened. Ex-type strains are denoted with “T”.

In section *Rubri*, four distinct *Monascus* species (*M. pilosus*, *M. purpureus*, *M. ruber*, and *M. sanguineus*) are recognized based on the well-supported phylogenetic lineage on the tree. To achieve higher phylogenetic resolution, an additional phylogenetic analysis was conducted using only section *Rubri* strains. The resulting tree revealed more clearly defined species boundaries among *M. pilosus*, *M. purpureus*, *M. ruber*, and *M. sanguineus* (**Fig. 3**).

**Figure 3.**
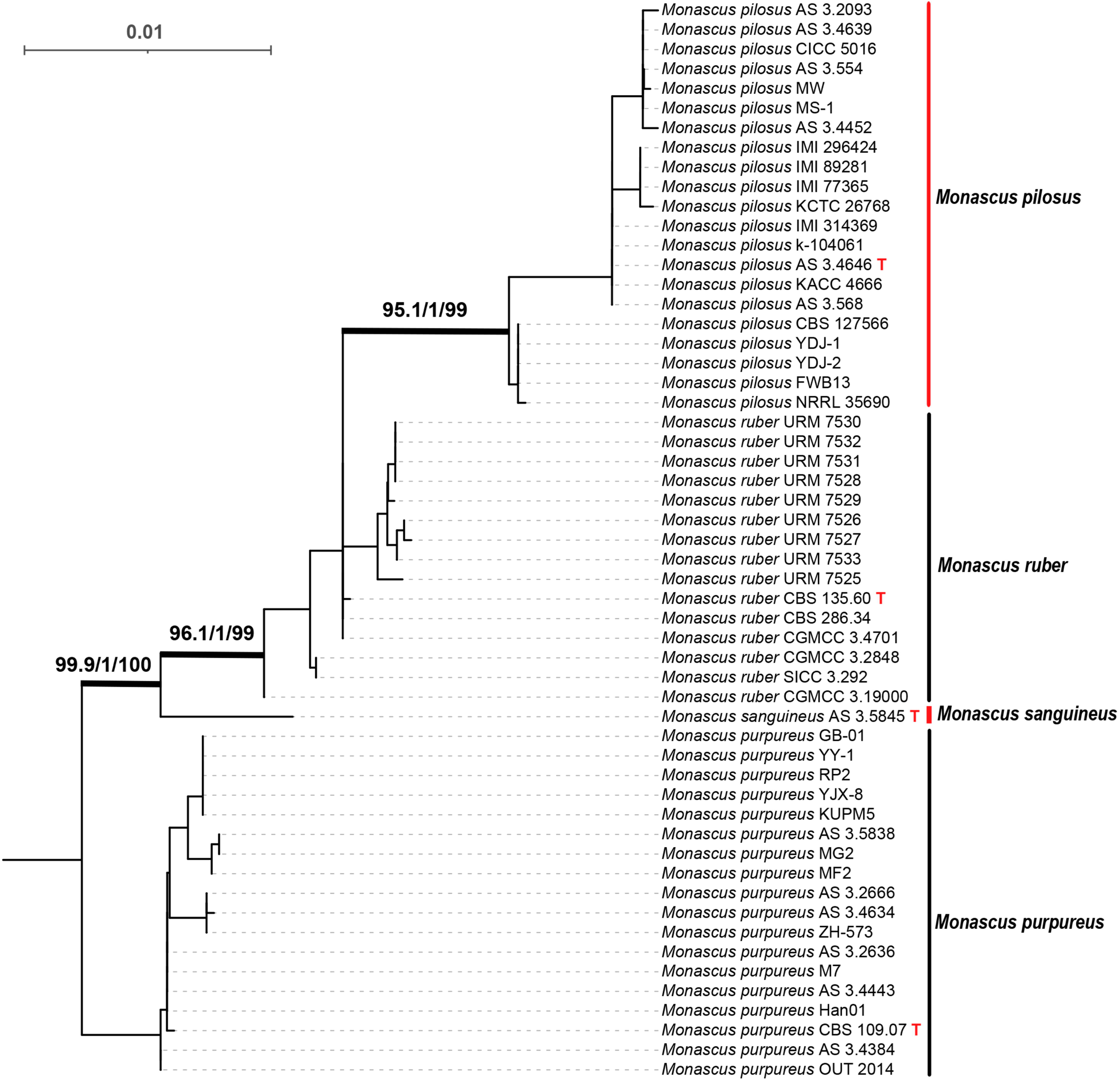
Concatenated phylogeny of the CaM, BenA, LSU, ITS and RPB2 gene regions illustrating the relationship within section Rubri. Numbers on the key branches indicate SH-aLRT support (%) / aBayes support / ultrafast bootstrap support (%). Ex-type strains are denoted with “T”.

Some species nomenclatures were revised on the basis of molecular phylogenetic analysis (**Figs 2**, **3**), with detailed information provided in Table 2. The analysis reveals that representative *M. pilosus* strains formed an independent monophyletic clade, which showed a clear boundary with the *M. ruber* clade, and thereby *M. pilosus* is not a synonym of *M. ruber*. The formerly recognized species *M. albidulus*, *M. anka*, *M. barkeri*, and *M. fumeus* were clustered within the *M. pilosus* clade and should therefore be treated as its synonyms. In addition, *M. purpureus* strains formed an independent monophyletic clade that exhibited a distinct boundary with the *M. sanguineus* clade, indicating that *M. sanguineus* is not a synonym of *M. purpureus*. The formerly recognized species *M. aurantiacus* and *M. rutilus* were clustered within the *M. purpureus* clade and should therefore be regarded as synonyms of *M. purpureus*.

### Taxonomy

By combination of previous studies (Barbosa et al. 2017; Visagie et al. 2025), the *Monascus* taxonomic system based on a curated DNA reference sequence database of *BenA*, *CaM*, *ITS, LSU,* and *RPB2* was updated by incorporating relevant data of synonymous species as well as the restored species *M. pilosus* and *M. sanguineus*. Nevertheless, to achieve reliable species delimitation within section *Rubri*, concatenated multi-locus datasets, rather than single-gene sequences, are recommended for molecular taxonomic analyses.

### 11 *Monascus* species are recognized and categorized into two sections ***Floridani*** and ***Rubri***

In *Monascus* section ***Floridani*** Barbosa & Houbraken (2017). MycoBank: #820076.

***M. argentinensis*** Stchigel & Guarro (2004). MycoBank: #500076. Typus: FMR 6778. Ex-type: CBS 109402 = FMR 6778.

DNA barcodes: *BenA* = KY709174; *CaM* = KY611935; *ITS* = JF922046; *LSU* = KY645974; *RPB2* = JN121423.

***M. flavipigmentosus*** Barbosa, Souza-Motta, Oliveira & Houbraken (2017). MycoBank: #820072.

Typus: URM 90064. Ex-type: CBS 142366 = URM7536 = DTO 353-A2.

DNA barcodes: *BenA* = KY709168; *CaM* = KY611929; *ITS* = KY511752; *LSU* = KY511782; *RPB2* = MN969201.

***M. floridanus*** Cannon & Barnard (1987). MycoBank: #132123.

Typus: IMI 282587. Ex-type: CBS 142228 = FLAS F54662 = CGMCC 3.5843 = BCRC33310 = UAMH 4180.

DNA barcodes: *BenA* = KY709172; *CaM* = KY611933; *ITS* = KY635848; *LSU* = KY635856; *RPB2* = KY611972.

***M. lunisporas*** Udagawa & Baba (1998). MycoBank: #446999.

Typus: SUM 3116. Ex-type: CBS 142230 = CGMCC 3.7951 = ATCC 204397 = NBRC33241 = BCRC 33640.

DNA barcodes: *BenA* = KY709171; *CaM* = KY611932; *ITS* = KY635847; *LSU* = KY635855; *RPB2* = KY611971.

***M. mellicola*** Barbosa, Souza-Motta, Oliveira & Houbraken (2017). MycoBank: #820073.

Typus: URM 90065. Ex-type: CBS 142364 = URM7510 = DTO 350-E6.

DNA barcodes: *BenA* = KY709143; *CaM* = KY611904; *ITS* = KY511726; *LSU* = KY511756; *RPB2* = KY611943.

***M. pallens*** Cannon, Abdullah & Abbas (1995). MycoBank: #413476.

Typus: IMI 356820. Ex-type: CBS 142229 = BSRA10266 = CGMCC 3.5844 = ATCC 200612 = BCRC 33641.

DNA barcodes: *BenA* = KY709173; *CaM* = KY611934; *ITS* = KY635849; *LSU* = KY635857; *RPB2* = KY611973.

***M. recifensis*** Barbosa, Souza-Motta, Oliveira & Houbraken (2017). MycoBank: #820074.

Typus: URM 90066. Ex-type: CBS 142365 = URM7524 = DTO 350-G6.

DNA barcodes: *BenA* = KY709157; *CaM* = KY611918; *ITS* = KY511740; *LSU* = KY511770; *RPB2* = KY611957.

In *Monascus* section ***Rubri*** Barbosa & Houbraken (2017). MycoBank: #820077.

***M. pilosus*** Sato ex Hawksw. & Pitt (1983). MycoBank: #109083.

*M. albidulus* Li & Guo (2004).

*M. anka* Nakaz. & K. Sato (1936) (nom. illegit.).

*M. barkeri* Dang. (1903).

*M. fumeus* Li & Guo (2004).

Typus: FRR 2194. Ex-type: IFO 4480 = AS 3.4646= ATCC 16363

DNA barcodes: *BenA* = PQ118070; *CaM* = PQ118100; *ITS* = PQ113596; *LSU* = PQ119540; *RPB2* = PQ118130.

***M. purpureus*** Went (1895). MycoBank: #235390.

*M. aurantiacus* Li ex Li & Guo (2004).

*M. rutilus* Li & Guo (2004).

Typus: IMI 210765. Ex-type: CBS 109.07 = IF04513 = ATCC 16426 = NRRL 1596 = FRR 1596.

DNA barcodes: *BenA* = KY709176; *CaM* = KY611937; *ITS* = KY635851; *LSU* = KY635859; *RPB2* = JN121422.

***M. ruber*** van Tieghem (1884). MycoBank: #234876.

Typus: IMI 81596. Ex-type: CBS 135.60 = IFO8451 = ATCC 15670.

DNA barcodes: *BenA* = KY709175; *CaM* = KY611936; *ITS* = KY635850; *LSU* = KY635858; *RPB2* = KY611974.

***M. sanguineus*** Cannon, Abdullah & Abbas (1995). MycoBank: #413477. Typus: IMI 356821. Ex-type: BSRA 10267 = AS 3.5845= ATCC 200613.

DNA barcodes: *BenA* = PQ118087; *CaM* = PQ118117; *ITS* = PQ113613; *LSU* = PQ119557; *RPB2* = PQ118147.

## Discussion

*Monascus* spp. are economically important filamentous fungi, and research on their classification is of great significance for their industrial applications. The genus *Monascus* belongs to the family Aspergillaceae, which is characterized by the production of stalked cleistothecial ascomata in the sexual stage and basipetospora in the asexual stage (Houbraken et al. 2020). Traditionally, the recognition of *Monascus* at the species level is based on their macro- and microscopic features, including the pigmentation of the cleistothecial walls, size and shape of conidia and ascospores, aerial hyphae, and growth rates on agar media (Barbosa et al. 2017). However, morphological taxonomy has its limitations. On the one hand, the identification of *Monascus* morphology may be affected by culture conditions such as medium composition, time, temperature, and humidity, as well as the subjective judgment of classifiers. For example, various color names have been used historically or are currently employed for the nomenclature of *Monascus* species. On the other hand, phenotypic characteristics are not necessarily indicative of the true genetic relatedness between taxa. For example, while cleistothecial features are key indicators in species recognition, no cleistothecia were observed in the species *M. mellicola* and *M. recifensis* (Barbosa et al. 2017). Therefore, the application of molecular typing to define species boundaries through unambiguous genetic markers, independent of culture conditions and subjective judgment, is likely to be a future trend in *Monascus* classification.

In this study, the phylogenetic relationships of 82 representative *Monascus* strains, encompassing 17 commonly accepted *Monascus* species, were studied using a concatenated dataset of five genes (*BenA*, *CaM*, *ITS*, *LSU*, and *RPB2*). Based on the findings of this study, 11 distinct *Monascus* species are recognized and categorized into two well-supported sections: *Floridani* (*M. argentinensis*, *M. flavipigmentosus*, *M. floridanus*, *M. lunisporas*, *M. mellicola*, *M. pallens*, and *M. recifensis*) and *Rubri* (*M. pilosus*, *M. purpureus*, *M. ruber*, and *M. sanguineus*). *M. pilosus* and *M. sanguineus* have been re-identified as separate species based on their well-supported and distinguishing phylogenetic lineages.

Species in section *Floridani* were isolated from natural habitats rather than traditional fermented products, while those in section *Rubri* are primary contributors to the production of related fermented foods. Generally, these two sections could also be delimited based on morphological characteristics (Barbosa et al. 2017). First, species in section *Floridani* exhibit restricted to no growth on media such as MEA, PDA, CYA, and G25N, whereas those in section *Rubri* show faster growth rates. Second, the production of *Monascus* pigments has not been reported in species from section *Floridani* to date, and their colony shades are typically light or brown, while those in section *Rubri* are renowned for their vivid *Monascus* pigments, with colony shades usually being orange, red, or dark brown. Third, fewer or no cleistothecia are observed in species from section *Floridani* compared to those in section *Rubri.* Furthermore, the taxonomic relationship within section *Floridani* is much more conserved and shares a broad consensus in the scientific community; however, that within section *Rubri* is the opposite, with quite a few controversies as mentioned below.

The data generated in the present study provide strong support for the recognition of *M. pilosus* as a distinct species. In a previous report, *M. pilosus* was regarded as a synonym of *M. ruber* (Barbosa et al. 2017). In the phylogenetic analysis conducted in that study, the authors used only a single strain, CBS 286.34, as the reference strain for *M. pilosus*. In contrast, in the phylogenetic analysis of the present study, strain CBS 286.34 was also clustered within the *M. ruber* clade, whereas other representative *M. pilosus* strains formed an independent monophyletic clade (**Figs 2, 3**). Consequently, strain CBS 286.34 is genetically affiliated with *M. ruber*, however, it was erroneously employed as a representative of *M. piolsus*, resulting in the conclusion that *M. pilosus* was a synonym of *M. ruber*. *M. pilosus* and *M. ruber* exhibit similar phenotypes, and they are frequently misidentified via traditional morphological classification. This is evidenced by the observation that some previously designated *M. ruber* strains were also placed within the *M. pilosus* clade (**Table 2**). In China, *M. pilosus* and *M. ruber* possess distinct industrial applications: *M. pilosus* is primarily used for the production of monacolin K, whereas *M. ruber* is mainly involved in the brewing of fermented foods. Additionally, *M. albidulus*, *M. anka*, *M. barkeri*, and *M. fumeus* are classified as synonyms of *M. pilosus* due to their close phylogenetic lineages (**Figs 2, 3**). Although these four species were isolated from fermented foods (**Table 2**), they have been rarely reported or rediscovered in previous studies.

Similarly, the data in the present study also support for the recognition of *M. sanguineus* as a distinct species. *M. sanguineus* was previously regarded as a synonym of *M. purpureus* (Barbosa et al. 2017). In the phylogenetic analysis of this study (**Figs 2, 3**), the ex-type strain of *M. sanguineus* (AS 3.5845) is situated between the *M. ruber*-clade and the *M. purpureus*-clade, maintaining a sufficient evolutionary distance from both the *M. purpureus* clade and the *M. ruber* clade. Furthermore, additional evidence, such as a compendium of biosynthetic gene clusters, indicates that *M. sanguineus* should be classified as a separate species from *M. purpureus* (He et al. 2018; Liu et al. 2022).

*M. ruber* and *M. purpureus* have consistently been regarded as distinct species since their discovery (**Table 1**). In the phylogenetic analysis of the present study, *M. ruber* strains did not form a fully independent monophyletic clade, whereas *M. purpureus* strains formed an independent monophyletic clade (**Figs 2, 3**). Additionally, *M. aurantiacus* and *M. rutilus* are considered synonyms of *M. purpureus* due to their close phylogenetic lineages (**Figs 2, 3**). These two species were isolated from fermented foods and named by Li and Guo (2004), yet they have rarely been mentioned or rediscovered in the existing literature.

In summary, traditional morphological approaches are insufficient for the discrimination of *Monascus* species, especially those with subtle morphological variations. This study proposes a molecular taxonomic approach for the genus *Monascus* based on the five-gene set (*BenA*, *CaM*, *ITS*, *LSU*, and *RPB2*). Generally, the phylogenetic resolution of these single genes is sufficient to differentiate *Monascus* species within section *Floridani*, whereas their concatenated multi-locus datasets should be used for accurate species delimitation in section *Rubri*. The genetic marker-based method proposed herein will substantially facilitate the accurate identification of *Monascus* species, thereby conferring considerable significance for related research and practical applications.

## Acknowledgments

The authors are very grateful to the Korean Collection for Type Cultures (KCTC) in South Korea for generously providing *Monascus* specimens and lab members Yanli Feng, Chaoyi Zheng and Qi Li for their efforts in strain collection.

## Funding

This work was financially supported by the National Natural Science Foundation of China (Nos. 31730068 and 31330059 to F. C.; No. 31601446 to W. C.).

## Competing interests

The authors have declared that no competing interests exist.

## Notes

### Competing Interest Statement

The authors have declared no competing interest.

